# Area- and Layer-Specific Organization of Timescales in Macaque Motor Cortex

**DOI:** 10.64898/2026.03.21.713374

**Authors:** Nilanjana Nandi, Laura López-Galdo, Simon Nougaret, Bjørg Elisabeth Kilavik

**Author notes:** co-senior authors (Simon Nougaret, Bjørg Elisabeth Kilavik).

## Abstract

Hierarchy in the brain emerges across spatial and temporal scales, enabling transformations from rapid sensory encoding to sustained cognitive control, and this organization is well established in sensory systems. In contrast, the hierarchical organization of the primate motor cortex remains debated, partly due to its agranular architecture and the absence of clear layer-constrained input-output projections. In particular, the relative hierarchical position of the dorsal premotor cortex (PMd) and the primary motor cortex (M1) cannot be resolved from anatomy alone. To investigate their relative organization, we adopted a unique multimodal approach using timescales derived from both single-unit spiking activity (SUA) and local field potentials (LFPs) in macaques performing a delayed-match-to-sample reaching task. We found convergent evidence for inter-areal temporal organization, with longer SUA timescales and smaller LFP aperiodic spectral exponents in M1. Across cortical depth, however, temporal dynamics depended on signal modality. LFP autocorrelation timescales were systematically longer in deep layers of M1, and this was accompanied by smaller LFP spectral exponents in deep layers in both areas. In contrast, SUA did not show significant laminar differences in timescales. Functionally, neurons with longer timescales exhibited more stable representations of the movement direction during movement preparation in PMd and broader temporal generalization during execution in both areas. Our results place M1 above PMd in the temporal hierarchy, and provide the first laminar characterization of SUA timescales in any cortical area. The divergence in laminar temporal organization between SUA and LFP possibly reflects their different physiological origins. Extracellular spikes capture neuronal output near the cell body, whereas LFPs primarily reflect synaptic population activity, potentially exhibiting layer differences in integration of apical and basal dendritic inputs.

**Highlights:** SUA and LFP-derived temporal dynamics place M1 above PMd in the cortical hierarchy

SUA shows no laminar differences in temporal dynamics

LFP exhibits significant layer-dependent temporal differences

Longer SUA timescales link to slower movement encoding dynamics

## Introduction

Hierarchy in the brain emerges across both spatial and temporal scales, enabling distributed yet coordinated processing of information across distinct regions. Cortical hierarchy is classically illustrated in the primate visual system (Felleman and Van Essen, 1991), where receptive fields expand in size and computational complexity from orientation-selective responses in V1 (Hubel and Wiesel, 1968) to abstract object representations in higher visual areas like inferotemporal cortex (Tanaka, 1996; DiCarlo, Zoccolan and Rust, 2012). Motor behavior likewise depends on multilevel transformations that convert sensory inputs into planned and executed actions. However, unlike sensory cortices, the organizational principles of the primate motor system remains less well understood, partly due to the agranular architecture of the motor cortex and the absence of clear layer-constrained input-output pathways that typically define feedforward and feedback projections (Shipp, 2005; Barbas and García-Cabezas, 2015).

The relative relationship between two motor areas involved in upper limb movements, the dorsal premotor cortex (PMd) and primary motor cortex (M1), has been debated. Early anatomical studies positioned M1 below PMd in the cortical hierarchy, interpreting PMd to M1 projections as descending feedback (Felleman and Van Essen, 1991; Shipp, 2005). Recent dual anterograde tract-tracing studies (Ninomiya *et al*., 2019) have shown layer-specific connectivity patterns between PMd and M1. In particular, PMd projections terminate predominantly in the superficial layers of M1, whereas M1 projections target both superficial and deep layers of PMd, suggesting a higher position for M1. These findings highlight the limitations of inferring motor hierarchy solely from anatomical connectivity.

Another line of work suggests that intrinsic temporal properties of neuronal activity may provide an alternative functional marker of cortical hierarchy (Murray *et al*., 2014). Intrinsic timescales refer to the temporal window over which neuronal activity integrates information and are typically estimated from the decay constant (τ or tau) of the autocorrelation of single-neuron spiking activity (SUA). Across the brain, timescales exhibit a hierarchical gradient. At the cortical level, they increased from sensory areas to higher-order regions like prefrontal cortex (Murray *et al*., 2014; Chaudhuri *et al*., 2015; Cavanagh, Hunt and Kennerley, 2020; Wang, 2021), whereas at the subcortical level, the input structures of the basal ganglia exhibited longer timescales than a downstream nucleus (Nougaret *et al*., 2021). Longer timescales support sustained information integration required for cognitive processes including working memory, decision-making, and motor planning (Cirillo *et al*., 2018; Wasmuht *et al*., 2018; Spitmaan *et al*., 2020; Golesorkhi *et al*., 2021; Manea *et al*., 2022; Zeraati *et al*., 2023; Trepka *et al*., 2024). Importantly such temporal organization across cortical areas has been shown across species (Zeisler *et al*., 2025), and using different signal modalities, including spiking activity (SUA), intracranial electroencephalography (iEEG) recordings (Cusinato *et al*., 2023), electrocorticography (ECoG) (Gao *et al*., 2020), and functional magnetic resonance imaging (fMRI) (Manea *et al*., 2022).

Timescales have not been systematically used to probe relative organization within the motor system (see, however (Morales-Gregorio *et al*., 2025)). Moreover, whether any laminar differences in temporal dynamics exist within a cortical area remains unknown. One theoretical proposal suggests that timescales should differ across layers, with longer timescales for neurons in superficial layers due to their enhanced recurrent connections (Cavanagh, Hunt and Kennerley, 2020). However, such laminar predictions have not been directly tested using single-neuron and LFP autocorrelations, both of which are direct measures of neuronal (population) timescales. A recent study (Halgren *et al*., 2021) reported that the LFP aperiodic spectral exponents (exp) decrease with cortical depth. This result was interpreted as evidence for longer timescales in superficial layers, based on the excitation/inhibition (E/I) framework of (Gao, Peterson and Voytek, 2017), in which larger spectral exponents indicate longer timescales. However, (Cusinato *et al*., 2023) directly examined the relationship between timescales and aperiodic spectral exponents in iEEG data, and reported that lower exponents were associated with longer autocorrelation-derived timescales across regions, consistent with an inverse relationship between exponent and timescales.

Here, we addressed three questions: (1) Do intrinsic timescales differ between M1 and PMd, and if so, what would these differences suggest about their relative hierarchical position? (2) Do timescales differ between superficial and deep layers? and (3) Do neurons with different timescales have different encoding properties during motor planning and execution? To answer these questions, we analyzed acute laminar recordings of SUA and bipolar LFPs from two male macaques performing a delayed-match-to-sample (DMS) arm-reaching task. First, we found that SUA-derived timescales were longer and LFP aperiodic exponents smaller in M1 than in PMd, suggesting a higher position for M1 in the cortical hierarchy. Second, LFP-derived measures exhibited systematic layer-dependent differences consistent with longer timescales in deep layers. In contrast, SUA timescales did not differ across layers. Third, neurons with longer tau exhibited more stable representations of movement direction during planning and execution.

## Results

We investigated intrinsic temporal dynamics in PMd and M1 of two macaques performing a DMS arm-reaching task (see Fig. 1A). To maintain comparable task structure across animals while reducing task demands, Monkey T performed sessions using a reduced set of three target positions, randomly selected on each session, whereas Monkey M performed the full set of four target positions. Neural activity (SUA and LFP) was recorded using acute laminar probes and single-tip electrodes (see Fig. 1B). For SUA, timescales (decay constant tau) were estimated using the autocorrelation method described by (Fontanier *et al*., 2022). Larger tau values indicate slower decay of temporal correlations and longer temporal integration (see Fig. 1C/D). For LFPs, bipolar derivations were computed across cortical depth to obtain local signal specificity in each channel (see Fig. 1E). While previous work has linked the aperiodic spectral knee to intrinsic timescales (Gao *et al*., 2020), reliable knee estimation was obtained in only ∼23% of channels in our dataset. We therefore adopted two complementary LFP-based approaches. First, LFP timescales were estimated from signal autocorrelation (ACF) using the timescales toolbox from Voytek and colleagues ((Cellier *et al*., 2025), <u>timescales toolbox</u>; see Fig. 1F - bottom). Second, aperiodic exponents have been proposed to reflect underlying E/I balance and have been linked to timescales (Gao, Peterson and Voytek, 2017; Halgren *et al*., 2021; Cusinato *et al*., 2023). Spectral exponents were estimated using FOOOF method (fitting oscillations and one over f, (Donoghue *et al*., 2020); fixed mode) in the 55-95 Hz range to capture the linear region of the spectrum, while avoiding contamination from knee frequencies (up to ∼45 Hz). This enabled us to obtain robust estimates of spectral decay (exp) across all channels (Fig 1F - top).

**Fig 1.**
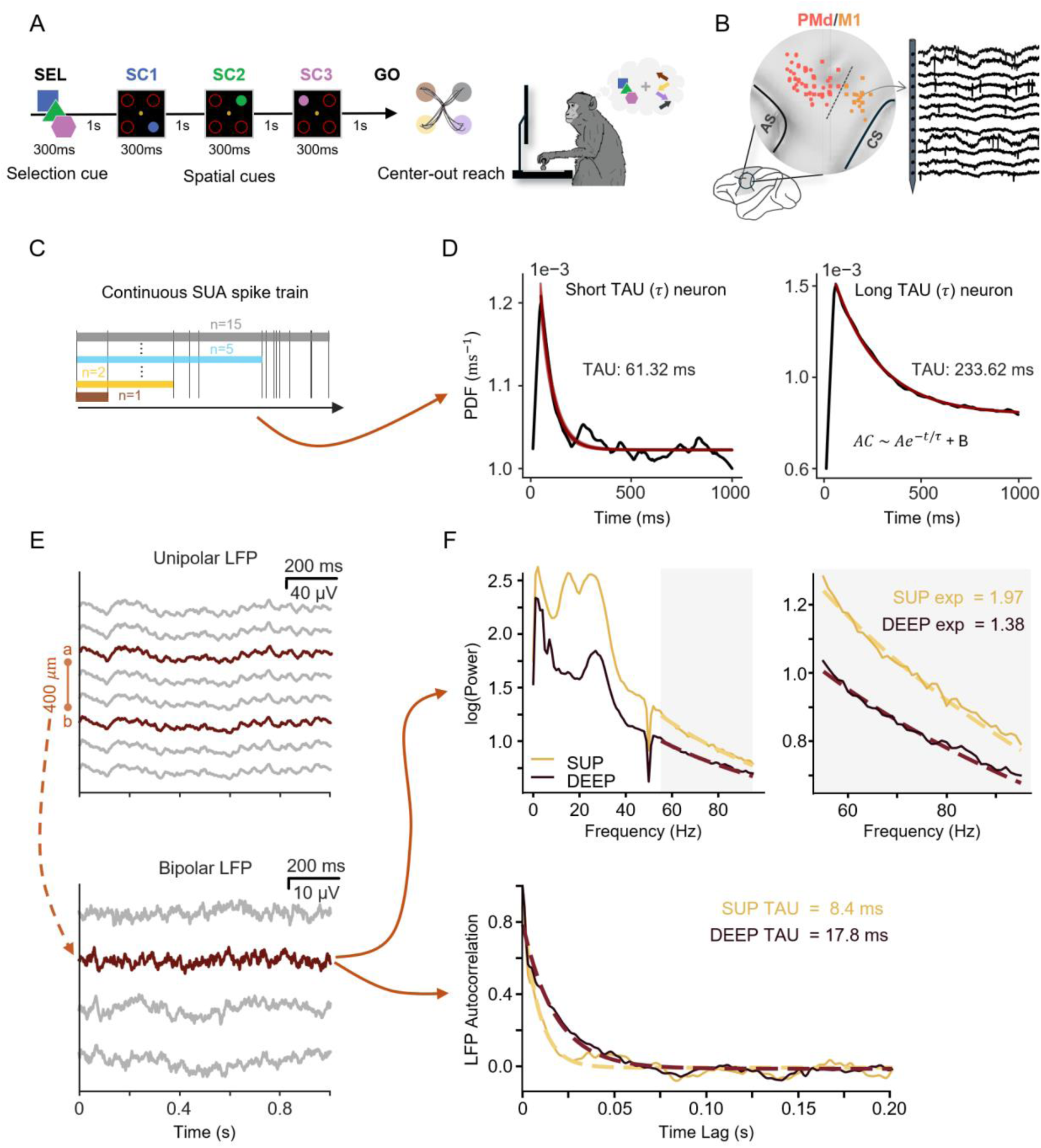
Experimental setting and methods to estimate temporal measures in SUA and LFPs. **(A)** Outline of the delay-match-to-sample (DMS) behavioral task and behavioral setup. In this task, the monkey must use a selection cue (SEL) to select the valid (color-matched) spatial cue (SC), to perform a centre-out movement in one out of four possible directions. Detailed description in the Methods section. **(B)** Schematic of recording sites in PMd and M1 from the two monkeys using acute laminar probes and single-electrodes (left), and example raw traces. (right) **(C)** Continuous spike train and computation of interspike interval (ISI) distributions across multiple orders (n=1, 2… 100). Vertical lines represent spikes from an example neuron. Colored bars (brown, yellow, blue, grey) correspond to multiple orders (n=1,2,5,15) across which interspike intervals (ISIs) are computed (up to n =100). **(D)** Spike timescale (tau/*τ*) estimation: The ISI distribution from (C) is used to compute a probability density function (PDF), from which the decay constant (tau) is estimated.The PDF of ISIs was fit with an exponential decay to extract tau. Example neurons with short (left) and long (right) tau are shown. **(E)** Example unipolar LFP channels (top) and corresponding bipolar derivations (bottom), computed by subtracting contacts (a-b) separated by 400 μm, and placing the new signal in the mid position between the two original contacts. **(F)** Light and dark colors indicate example superficial (SUP) and deep (DEEP) bipolar LFP channels, respectively. (Top) Aperiodic exponents were estimated by fitting power spectral densities using the FOOOF algorithm in the 55–95 Hz range. (Bottom) LFP timescales were extracted by fitting an exponential function to the autocorrelation function (ACF) of the bipolar LFP signal using timescales toolbox.

### Cortical organization based on temporal dynamics between M1 and PMd

Recording sites were classified as PMd or M1 based on a functional categorization. In a recent study using this dataset (Nougaret *et al*., 2024), we described that a low beta band rhythm is dominant in posterior (M1) LFPs and reflects movement preparation and postural dynamics, while a high beta band rhythm is dominant in anterior (PMd) LFPs and reflects spatiotemporal attention. This clear functional organization was used to delineate a functional border between M1 and PMd at the transition between low and high beta dominance (see Fig. 1B).

Timescales have been proposed as a marker of cortical hierarchy across brain areas. To test whether such organization is present within the motor cortex, we compared spike and LFP timescales (tau) and LFP aperiodic spectral structure (exp) between these functionally defined PMd and M1 areas. SUA timescales were longer in M1 than in PMd, see Fig. 2A (top). All statistical results are shown in Table 1. At the level of individual monkeys, this difference also reached significance in monkey M (see Supplementary Table 1). In contrast, LFP autocorrelation timescales did not differ between M1 and PMd (Fig. 2B (top)), though a marginal difference emerged if considering only distinct task epochs (preGo epoch, see control analyses below). However, LFP aperiodic spectral exponents were significantly lower in M1 than in PMd (Fig. 2C, top; Table 1), an effect that was also significant when quantified individually in monkey M (see Supplementary Table 1). This inverse relation between aperiodic exponents and SUA timescales is consistent with previous findings (Cusinato *et al*., 2023).

**Fig 2.**
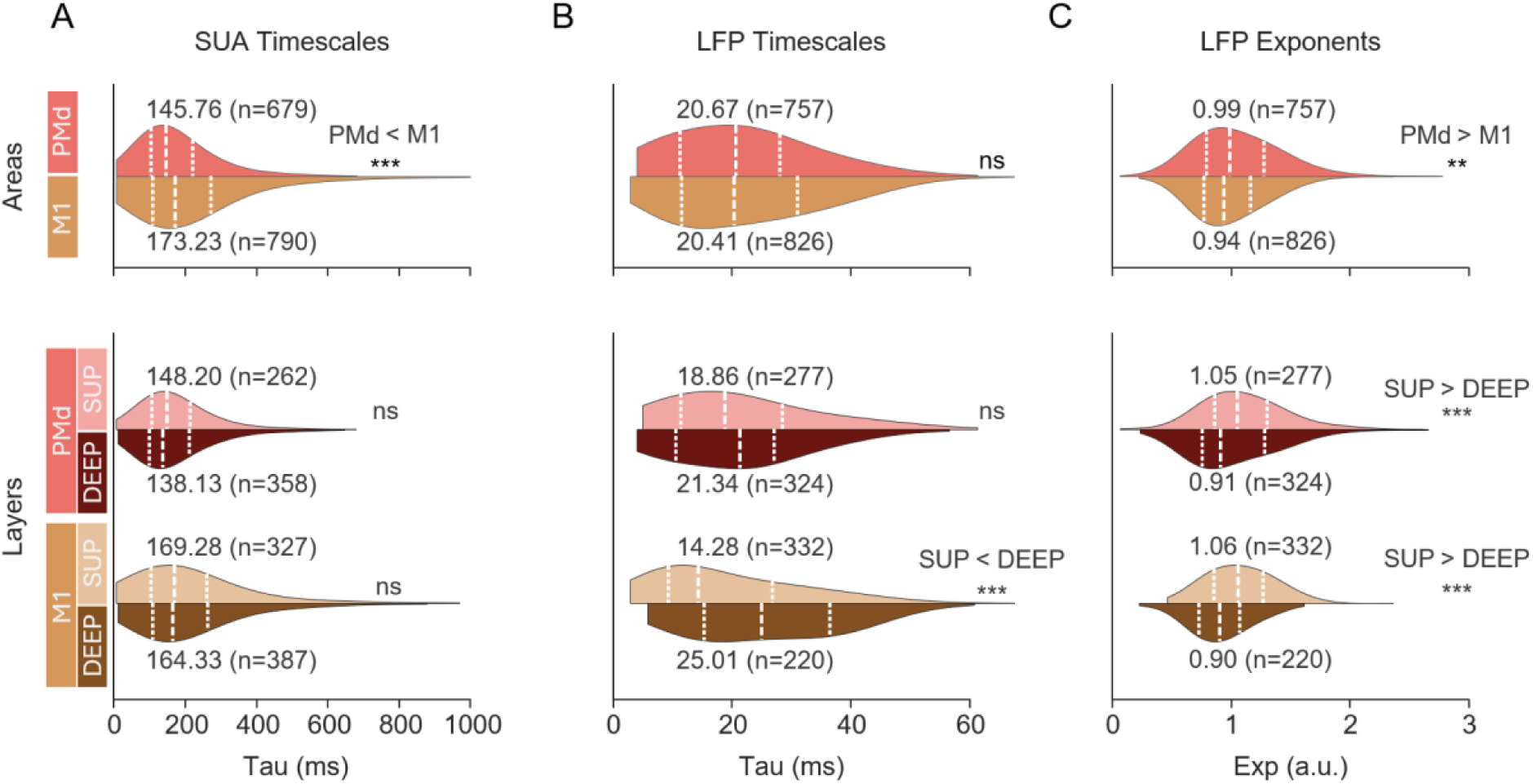
Characterization of spike and LFP temporal measures across cortical areas and layers. Violin plots showing the distributions of SUA-derived timescales **(tau; A)**, LFP-derived timescales **(tau; B)**, and LFP aperiodic spectral exponents **(exp; C)**. Top panels compare recordings from PMd and M1, whereas bottom panels compare superficial (SUP) and deep (DEEP) layers within each area. Lighter shades indicate superficial layers and darker shades indicate deep layers. Dashed lines indicate the median, 1st and 3rd quartiles values for each area. Numbers indicate median values, and n denotes the number of neurons **(A)** or bipolar LFP channels **(B–C)**. Statistical comparisons were performed using Kruskal-Wallis tests. Asterisks denote significant differences (**p < 0.05, ***p < 0.0005); ns, not significant.

**Table 1.** Statistical comparisons of tau and exp (reported median values)

| Comparison by area / layer |  |  | Kruskal Wallis |
| --- | --- | --- | --- |
|  | PMd | M1 |  |
| **Spike tau** | 145.76ms (n=679) | 173.23ms (n=790) | $\chi^2=13.7$, $p=0.0002$ |
| **LFP tau** | 20.67ms (n=757) | 20.41ms (n=826) | $\chi^2=2.91$, $p=0.09$ |
| **LFP exp** | 0.99 (n=757) | 0.94 (n=826) | $\chi^2=10.77$, $p=0.001$ |
|  | PMd SUP | PMd DEEP |  |
| **Spike tau** | 148.2ms (n=262) | 138.13ms (n=358) | $\chi^2=1.1$, $p=0.3$ |
| **LFP tau** | 18.86ms (n=277) | 21.34ms (n=324) | $\chi^2=0.2$, $p=0.64$ |
| **LFP exp** | 1.05 (n=277) | 0.91 (n=324) | $\chi^2=14.58$, $p=0.0001$ |
|  | M1 SUP | M1 DEEP |  |
| **Spike tau** | 169.28ms (n=327) | 164.33ms (n=387) | $\chi^2=2.75$, $p=0.1$ |
| **LFP tau** | 14.28ms (n=332) | 25.01ms (n=220) | $\chi^2=47.6$, $p=5.34e-12$ |
| **LFP exp** | 1.06 (n=332) | 0.9 (n=220) | $\chi^2=42.6$, $p=6.75e-11$ |

To determine whether these PMd-M1 differences depended on task-related neural activity, we performed several control analyses. Previous studies have shown that although timescales can vary with behavioral demands, their relative ordering across cortical regions remains stable (Manea *et al*., 2023). We therefore estimated timescales from two task epochs: preSC1, before directional information was provided with the cue, and preGo, while directional information was actively being maintained in working memory (see Materials and Methods). We additionally utilised the SUA to estimate tau from the full-trial epoch in correct trials only, thus excluding error trials, longer pauses in behavior and the intertrial intervals. Control analyses confirmed that both spike- and LFP-derived measures across task epochs were correlated (all p < 0.0001; see Supplementary Fig. 1D-F and Supplementary Fig 2C-D), supporting the robustness of these measures. Consistent with the main analysis, the relative PMd/M1 temporal organization was preserved across epochs, with M1 generally exhibiting longer timescales and lower aperiodic exponents than PMd. These differences reached significance for spike-derived tau during the preSC1 baseline epoch (p = 0.01), for LFP-tau during the preGo epoch (p = 0.03), and for aperiodic exponents across all epoch comparisons (all p < 0.01; Supplementary Fig. 1A-C; Supplementary Fig. 2A-B).

Together, these results indicate a temporal gradient across the motor cortex, with longer SUA timescales and flatter aperiodic exponents in M1 relative to PMd, that is robust across task epochs.

### Laminar organization of temporal dynamics is specific to LFPs, not spikes

Having established a temporal organization across M1 and PMd, we next examined whether similar gradients were present across cortical depth. Motivated by prior hypotheses proposing the existence of laminar differences in temporal integration (Cavanagh, Hunt and Kennerley, 2020) and anecdotic experimental observations in LFP aperiodic spectral structure (Halgren *et al*., 2021), we compared SUA- and LFP-derived temporal metrics between superficial and deep layers. Units and channels were classified as superficial (SUP) or deep (DEEP) based on relative cortical depth (see Materials and Methods).

For SUA, neither of the two areas had significant differences in tau across layers (Fig. 2A (bottom)). For individual monkeys, only deep layers in M1 showed longer timescales compared to superficial layers in monkey M, but this was not confirmed in PMd, nor in any of the two areas for monkey T (see Table 1; Supplementary Table 1).

LFP timescales were longer in deep than superficial layers in M1 but a similar tendency in PMd did not reach significance (see Fig. 2B (bottom)). In individual animals, deep layers in M1 exhibited significantly longer timescales compared to superficial layers in M1 in both monkeys, and in PMd in monkey M (see Table 1; Supplementary Table 1). Furthermore, the LFP aperiodic exponents were smaller in deep than superficial layers in both PMd and M1 (see Fig. 2C (bottom); Table 1). This effect was also significant in individual animals in PMd in monkey M and in M1 in Monkey T (see Supplementary Table 1).

Together, these results suggest that laminar temporal organization is more consistently reflected in LFP signals than in spiking activity.

### Neural timescales shape movement-direction coding dynamics

Neurons with longer timescales have been shown to maintain more stable and sustained representations of task-relevant information, supporting persistent coding during working memory (Wasmuht *et al*., 2018; Cavanagh, Hunt and Kennerley, 2020). (Cirillo *et al*., 2018) showed stronger directional information in long-tau neurons during movement planning in PMd. However, whether this timescale-dependent directional selectivity reflects a more persistent encoding, and whether it can also be observed during movement execution in PMd and in M1, remains to be established. We therefore examined how timescales relate to movement direction encoding across the planning and execution periods of our task. First, we focused on a period from the appearance of the selection cue (SEL) to the GO signal. Monkeys knew from the onset of the valid SC (SC1 in blue trials, SC2 in green trials and SC3 in pink trials) the future direction of the movement they had to perform after the GO signal. PMd and M1 neurons exhibited distinct profiles of movement direction encoding (Fig. 3A). As a population, PMd neurons encoded movement direction after the occurrence of the valid SC in each color condition, peaking transiently at about 80 to 90% decoding accuracy, then decreasing and stabilizing movement direction information until the appearance of SC3 (for blue and green trials), before ramping up until after the GO signal to reach its maximum around movement onset (Fig. 3A, top). In contrast, M1 neurons encoded movement direction only from about 500ms prior to the onset of the GO signal, independently of color condition, reaching maximum decoding accuracy around movement onset (Fig. 3A, bottom).

**Fig 3.**
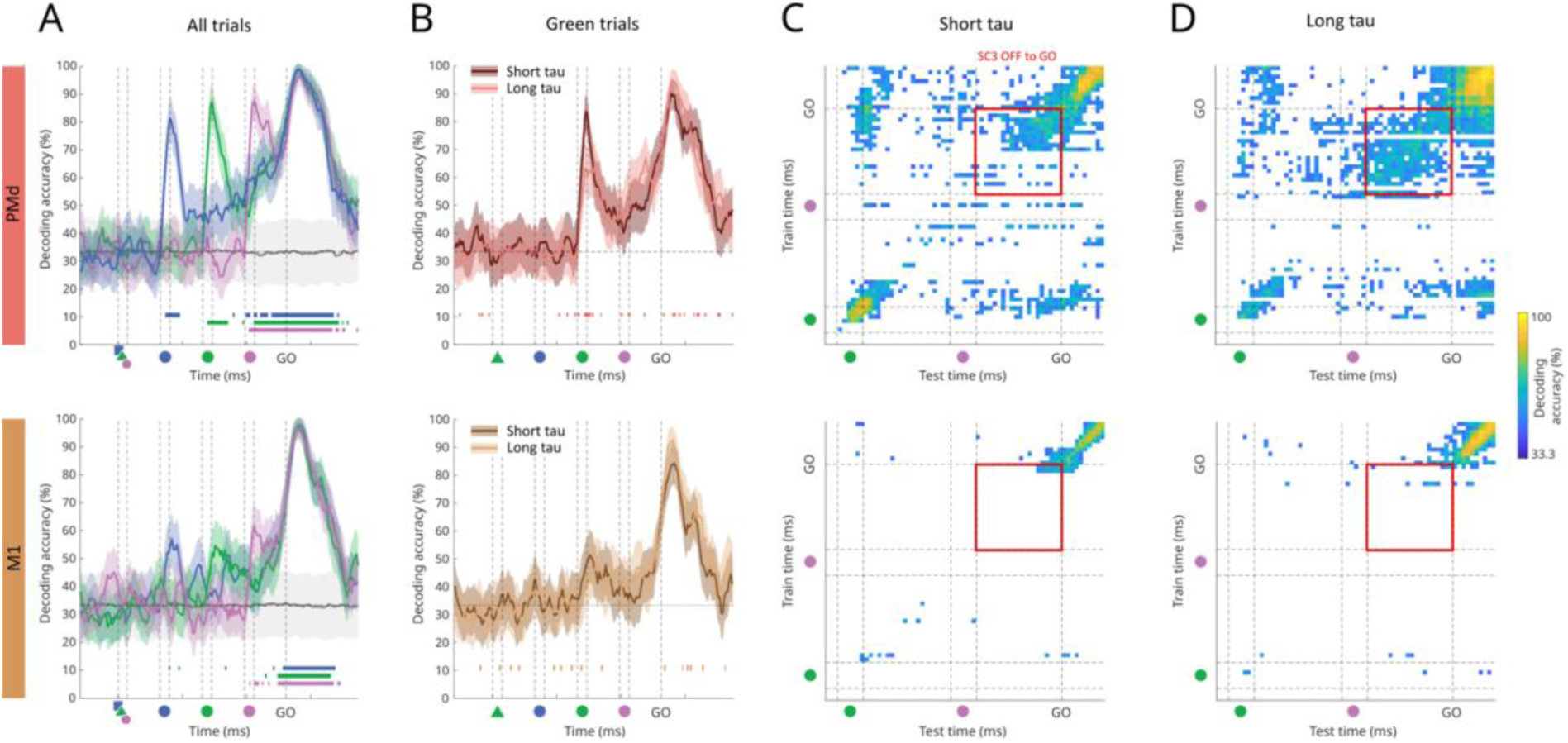
Direction encoding in short and long tau neurons during movement planning. PMd at the top, M1 at the bottom. **(A)** Time-resolved classification accuracy (mean ± SD) of a linear classifier trained to discriminate between 3 different movement directions (chance level = 33%) based on population spiking activity. Accuracies are computed separately for the three color conditions, blue, green and pink. Gray curve represents the accuracies when labels have been shuffled (for readability, only one shuffle curve is plotted). Coloured tick marks below the curves indicate time bins with classification significantly above chance level (Monte-Carlo p-value approximation < 0.05). Decoding performance has been smoothed on 5 successive bins for display purposes. **(B)** Similar representation, only for green trials, for decoding performance of two classifiers built with either the 50% of short or long tau neurons subpopulations. Coloured tick marks below the curves indicate time bins with significant difference between the decoding performance of either subpopulation (mean higher-accuracy condition > 97.5th percentile of lower-accuracy condition distribution). **(C)** Cross-temporal decoding maps for short-tau neuronal population, in green trials. Only significant pixels are shown in colour, i.e. higher decoding accuracy in the original data than the 95th percentile of accuracy of surrogate data with randomly shuffled movement direction labels. Off-diagonal pixels were only considered when decoding at the corresponding training time was significant, ensuring that temporal generalization was assessed only from points showing reliable decoding. The colour of each pixel represents the decoding accuracy in the original data. The red square represents the area in which the significant pixels are counted for statistical comparison across short and long tau subpopulations, taken in the 1 sec delay before GO. **(D)** Same as in C, for long tau neuronal population.

First we asked whether timescales influence the instantaneous encoding strength of task-relevant information. As illustrated for green trials in Fig. 3B, apart from a few isolated bins, overall the decoding profile was largely comparable between short- and long-tau neuronal populations in PMd and M1. We only observed, in PMd neurons after SC2 (Figure 3B, top) and in M1 neurons after SC3 (Supplementary Figure 3D), a higher peak of decoding in short tau neurons for 200ms and 250ms respectively. These results indicate that timescales do not substantially alter the amount of encoded information. We next examined whether timescales instead affect the temporal stability of neural representations using cross-temporal decoding during the final delay epoch (SC3_OFF to GO), when decoding accuracy of movement direction was significantly above chance across all (SEL) color conditions. Long-tau PMd neurons supported markedly more stable direction representations than short-tau neurons, as reflected by a greater proportion of significant cross-temporal decoding pixels (green SEL : 144/400 [36.00%] vs 267/400 [66.75%] permutation test, p < 0.0001; Fig. 3C–D top). The same effect was observed in pink trials (p <0.0001, Supplementary Fig 3D) and the same trend observed in blue trials (p=0.15, Supplementary Fig 3A). In contrast, M1 neurons showed very few significant pixels overall during this period, regardless of tau group (green: 12/400 [3.00%] vs 9/400 [2.25%] ; p = 0.66; Fig. 3C–D bottom; Supplementary Fig. 3), largely explained by the absence of instantaneous decoding.

We further examined the decoding profile of short and long-tau populations during movement execution. In both areas, movement direction was equally well decoded by short and long tau neuronal activity, in terms of temporal profile and accuracy (Fig. 4A). The accuracy increased before movement onset, reached its maximum, close to 100% between 150 and 300 ms after movement onset and slowly decreased after movement completion. However, the cross-temporal decoding profiles of short and long tau populations showed striking differences in the width of the decoding diagonal (Fig. 4B, C). The diagonals were broader for long tau compared to short-tau neurons. To quantify this difference, decoding accuracies from train times 0 to 500ms (see red parallelogram in Fig. 4B top/left) were realigned to obtain a distribution of values centered at lag 0. The central value of this distribution is the average of the diagonal values from 0 to 500ms.

**Fig 4.**
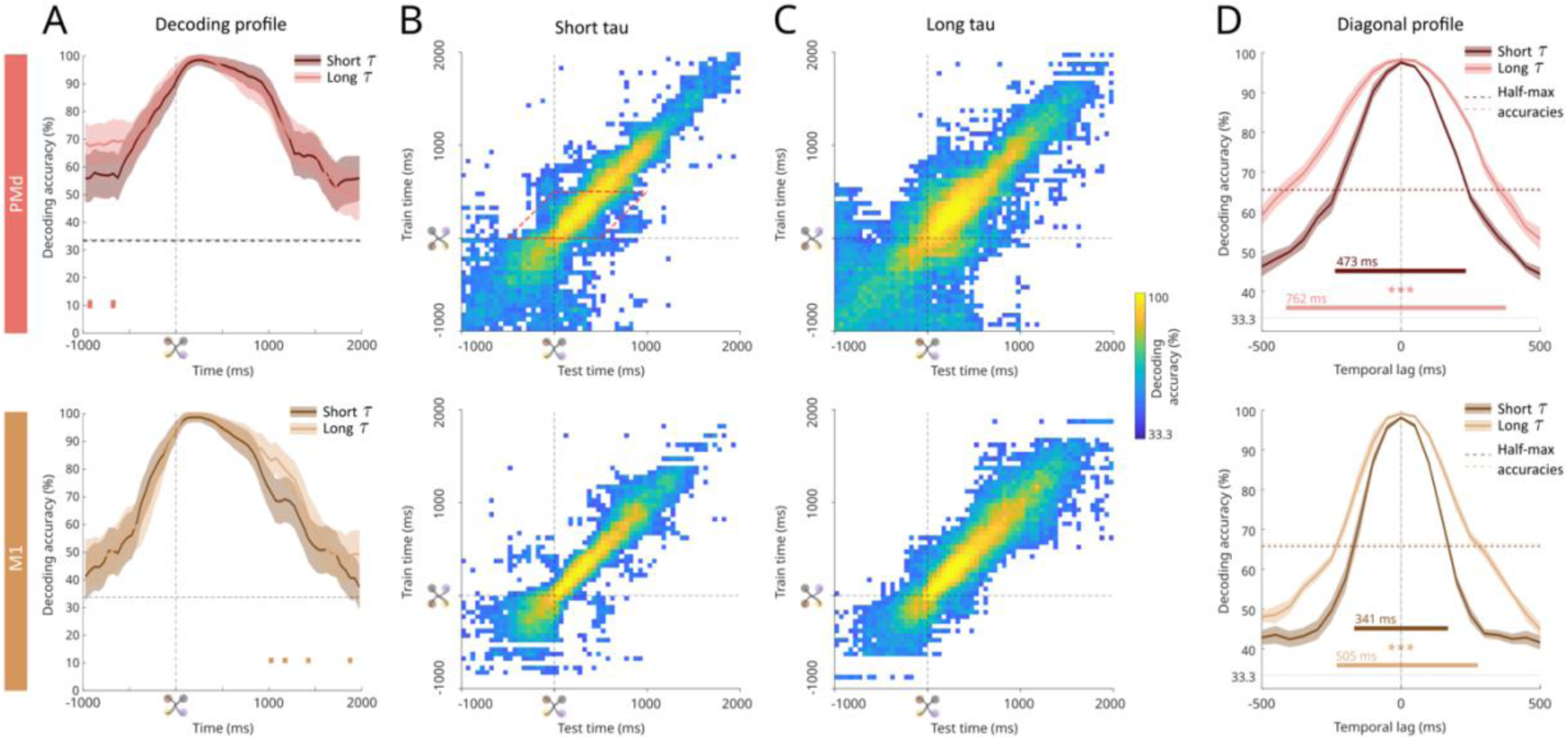
Direction encoding in short and long tau neurons during movement execution. PMd at the top and M1 at the bottom. **(A)** Time-resolved classification accuracy (mean ± SD) of a linear classifier trained to discriminate between 3 different movement directions (chance level = 33%) based on short or long tau population spiking activity, for the three-color conditions combined. Coloured tick marks below the curves indicate time bins with significant difference between the two (mean higher-accuracy condition > 97.5th percentile of lower-accuracy condition distribution). **(B)** Cross-temporal decoding maps for short tau neuronal population, from -1000ms to +2000ms around movement onset. Only significant pixels are shown in colour, i.e. higher decoding accuracy in the original data than the 95th percentile of accuracy of surrogate data with randomly shuffled movement direction labels. Off-diagonal pixels were only considered when decoding at the corresponding training time was significant, ensuring that temporal generalization was assessed only from reliable decoding models. The colour of each pixel represents the decoding accuracy in the original data. The red parallelogram illustrates the epoch from which the diagonal profiles were computed. **(C)** Same as in C, for long tau neuronal population. **(D)** Diagonal profile and half-max accuracies averaged across the first 500ms after movement onset. ***, permutation test, p<0.0005.

Based on this value, for each population of neurons, the half max windows were computed. For both M1 and PMd neurons, the diagonal profiles were systematically wider in long-tau neurons (permutation test, p<0.001) (Fig. 4D). A complementary analysis with neurons separated into quartiles instead of being classified based on the median tau, confirmed the association between larger tau values and wider diagonal (Supplementary Figure 4).

## Discussion

Here, using a multimodal approach combining spiking activity and LFPs, we investigated the hierarchical organization of motor cortical areas and layers through the lens of timescales. We report three main findings. First, both neural timescales and LFP spectral exponents suggested that M1 is placed above PMd in the temporal hierarchy. Second, at laminar resolution, LFP-derived measures showed consistent layer-dependent differences. Spectral exponents were significantly smaller in deep than superficial layers in both M1 and PMd, while LFP-derived autocorrelation timescales were longer in deep layers in M1. In contrast, spiking activity did not show any laminar differences in temporal dynamics. Third, probing the functional relevance, we found that neurons with longer tau exhibited more stable representations of memorized movement direction during the planning phase in PMd and during execution in both areas.

Timescales have been proposed as an intrinsic organizational principle of the brain. Prior work has shown that estimation of tau from different signal types often converge on similar hierarchical ordering across cortical areas. Our simultaneous observations across SUA and LFP, in two areas and layers, suggest that tau from different signals captures distinct aspects of neural population activity. Neural timescales derived from single unit activity reflect the duration over which the firing activity of individual neurons remains correlated over time. The timescales obtained from autocorrelation of LFP signals capture the temporal persistence of population-level activity, where longer timescales reflect more sustained, slow-varying fluctuations within the local circuit. Aperiodic exponents quantify the slope of scale-free activity, which has been proposed to reflect, amongst other factors, the excitation/inhibition (E/I) balance within local populations. Larger exponents correspond to a steeper suppression of higher frequency power. The inverse relation observed between timescales and aperiodic exponents suggests that the mechanisms shaping scale-free spectral structure may be different from those supporting sustained autocorrelated population activity. Stronger suppression of high-frequency activity may not necessarily translate into longer temporal persistence of the LFP signal, resulting instead in faster decorrelation and shorter autocorrelation timescales.

Together, these three measures provided complementary windows into the temporal organization of the motor cortex, both across areas and within the laminar microcircuit.

### Cortical hierarchy between PMd and M1 in light of temporal dynamics

The concept of anatomical hierarchy is well established in sensory systems, where laminar projection patterns define the direction of information flow. Feedforward projections terminate in layer IV to provide driving input to higher-order areas, whereas feedback projections avoid layer IV and exert modulatory influences on lower areas. In contrast, the motor cortex lacks a distinct granular layer IV, complicating hierarchical inference based solely on laminar projections. Functionally, M1 appears to occupy the terminal stage of the motor hierarchy, as it provides the densest corticospinal output and low-intensity stimulation readily evokes movements, which would predict a feedforward projection from PMd to M1. However, available anatomical evidence presents a paradox - projections from PMd to M1 display feedback-like termination patterns, suggesting a modulatory rather than purely driving influence (Shipp, 2005). A more recent dual anterograde tracing study (Ninomiya *et al*., 2019) demonstrated that PMd projections terminate primarily in deeper portions of L2/3 and weakly in layer 1 of M1, while M1 sends projections back to superficial and deep layers of PMd, supporting an ascending influence from PMd to M1 and placing M1 higher in the motor cortical hierarchy. Similarly (Barbas and García-Cabezas, 2015) proposed that M1 is the most structurally specialized motor area, exhibiting extensive myelination, restricted long-range connections, and elaborate cortical architecture, whereas premotor areas exhibit an intermediate structural gradient toward M1. Together, these findings highlight the difficulty of resolving motor hierarchy using classical anatomical criteria alone and motivate the use of functional metrics such as timescales.

Timescales that quantify the temporal window over which neurons sustain information, have been shown to correlate with hierarchical organization across cortex (Murray *et al*., 2014). Using this temporal framework, we found that M1 neurons exhibit longer timescales than PMd, supporting its hierarchical placement above PMd, consistent with anatomical evidence from anterograde tracing studies (Ninomiya *et al*., 2019). At the population level, timescales have also been estimated from LFP signals (Gao *et al*., 2020). Although identifying a knee frequency in the power spectrum was feasible in only a subset of our channels, alternative estimates based on LFP autocorrelation and spectral decay yielded complementary measures of temporal dynamics across areas. Aperiodic spectral exponents were significantly smaller in M1 than PMd. This pattern aligns with recent intracranial iEEG findings showing that spectral exponents flatten as timescales increase along sensory hierarchies (Cusinato *et al*., 2023), thus corroborating a higher position in the hierarchy for M1 compared to PMd. However, LFP autocorrelation timescales did not differ significantly between the two areas, suggesting that spectral scaling and temporal persistence of population fluctuations may reflect dissociable aspects of local circuit dynamics.

Regardless of this dissociation, a key question is whether the difference in temporal organization observed between PMd and M1 reflects intrinsic properties or is caused by task demands. To address this, we estimated neural timescales and LFP measures during specific task epochs - preSC1, when no directional information was available and working memory was not engaged, and preGo, when directional information had to be actively maintained in working memory during motor planning. Despite the markedly different cognitive demands of these epochs, both SUA timescales and LFP-derived measures remained highly correlated across epochs (p < 0.0001; Supplementary Figs 1D-F & 2C-D), supporting their intrinsic nature independent of task engagement. Furthermore, consistent with previous reports (Manea *et al*., 2024), whereas the absolute values of these measures differed across epochs, the relative ordering between PMd and M1 remained stable. The observed expansion in SUA timescale magnitude across preSC1/preGo epochs (∼80 ms increase in median) was not accounted for by the removal of inter-trial intervals, as estimating tau from the complete trial did not produce a comparable increase (see Supplementary Fig. 1C), suggesting it may be inherent to the autocorrelation estimation method itself. We used the approach of (Fontanier *et al*., 2022), which has been shown to yield timescale values correlated with those from the more traditional method of (Murray *et al*., 2014), while permitting inclusion of a larger proportion of neurons.

Together, these findings suggest that the temporal organization between PMd and M1 reflects stable intrinsic properties of these areas rather than transient task-driven states, motivating the question of whether similar temporal gradients extend to the laminar microcircuit within each area.

### Modality-dependent laminar organization of temporal dynamics

Neural timescales exhibit substantial heterogeneity within cortical areas, reflecting diversity in local microcircuit dynamics. Such variability is thought to support flexible computation, allowing neural populations to simultaneously process fast-changing inputs and integrate information over longer temporal windows (Wasmuht *et al*., 2018; Cavanagh, Hunt and Kennerley, 2020; Stern, Istrate and Mazzucato, 2023). Theoretically, part of this diversity has been proposed to be expressed also across layers, with longer timescales for neurons in superficial layers due to their enhanced recurrent connections (Cavanagh, Hunt and Kennerley, 2020). However, this prediction about timescale differences across layers has not been directly tested in any cortical area.

In the motor cortex, this question is further complicated by the absence of a clearly defined layer IV, limiting classical interpretations of laminar specialization. Some evidence points to laminar differentiation: for example, superficial PMd neurons have been reported to carry choice-related signals earlier and more prominently than deeper neurons (Chandrasekaran *et al*., 2017). At the same time, other studies emphasize coordinated processing across layers within local cortical columns rather than strict laminar segregation. Recordings in PMd (Opris *et al*., 2011; Opris, 2013) introduced the concept of functional ‘minicolumns’, demonstrating stronger synchronization between superficial (L2/3) and deep (L5) neurons within the same column than across adjacent columns. More recently, another study (López-Galdo *et al*., 2025) proposed that adaptive computations during sensorimotor transformation and movement planning emerge from coordinated activity spanning the entire motor cortical column, rather than from discrete layer-specific processing. In this framework, the column acts as an integrated computational unit, with dynamic fluctuations in laminar contributions shaping choice- and movement-related representations over time. Consistent with this framework, we did not observe significant laminar differences in SUA timescales in either PMd or M1. Instead, neurons with both short and long timescales were found in both superficial and deep layers, indicating substantial within-layer heterogeneity but little evidence for systematic laminar specialization of temporal dynamics.

In contrast, LFP-derived measures exhibited systematic laminar differences. LFP autocorrelation timescales were significantly longer in deep compared to superficial layers in M1. This laminar effect in M1 was evident both at the group level and in individual animals. A similar trend was observed in PMd at the group level, reaching significance in one animal. This dissociation between spiking and LFP measures suggests that laminar differences in temporal dynamics may preferentially emerge in mesoscopic signals, potentially reflecting layer-specific synaptic and dendritic dynamics that are not readily captured at the level of action potentials of individual neurons.

LFP-derived aperiodic exponents were furthermore systematically smaller in deep relative to superficial layers in both M1 and PMd. The decrease in the aperiodic (1/f) slope with cortical depth has been reported previously (Halgren *et al*., 2021). In that study, smaller slopes in deep layers were interpreted as reflecting shorter timescales, based on a proposed theoretical framework (Gao, Peterson and Voytek, 2017; Ameen *et al*., 2025). In that framework, stronger inhibitory activity, associated with slower, longer-lasting synaptic currents, is predicted to increase the temporal autocorrelation structure of population activity (longer timescales), resulting in a steeper spectral slope (larger exponent). Conversely, relatively stronger excitation is associated with flatter spectra (smaller exponent) and reduced temporal correlations (i.e. shorter timescales). However, (Halgren *et al*., 2021) did not directly estimate timescales from LFP autocorrelation, leaving this interpretation untested. Our direct measurements stand in contrast to this interpretation. Smaller exponents in deep layers co-occurred with longer LFP autocorrelation timescales, consistent with the empirical finding that aperiodic exponents inversely mirror autocorrelation-based timescales across brain regions (Cusinato *et al*., 2023). Together, these LFP-derived measures converge on the interpretation that deep layers exhibit longer temporal integration at the population level. These findings diverged from theoretical predictions of longer timescales in superficial layers (Cavanagh, Hunt and Kennerley, 2020), and suggest that spectral slope alone may not provide a reliable proxy for temporal persistence. To our knowledge, this is the first study to directly estimate timescales across cortical layers using multiple complementary signals and measures.

The observed dissociation between spike- and LFP-derived measures across layers may reflect not only the distinct neural signals they capture, but also the particular geometry of the laminar microcircuit. Spiking activity captures the localised output of individual neurons at the recording site, whereas LFP signals primarily arise from synaptic inputs and dendritic processing, largely dominated by excitatory post-synaptic potentials. In layers L5/L6, pyramidal neurons extend apical dendrites toward superficial layers, spanning much of the cortical column. Bipolar LFP signals at a particular depth will integrate synaptic inputs predominately at that depth, possibly highlighting laminar differences in temporal dynamics of apical/distal and basal/proximal dendrites at the population level that remain invisible at the level of individual neuron spiking outputs.

### Timescales shape temporal dynamics of movement direction encoding during planning and execution

Timescales are believed to structure the temporal dynamics of neural coding, with long-tau neurons supporting stable representations and short-tau neurons supporting more transient ones. Evidence from the prefrontal cortex links longer timescales to sustained working memory representations (Murray *et al*., 2014; Cavanagh *et al*., 2016, 2018), whereas (Cirillo *et al*., 2018) reported that longer-tau neurons in PMd showed stronger response selectivity during a delay period in which monkeys were preparing their future spatial response. In our study, we examined two motor cortical areas, PMd and M1, which are associated with movement planning and execution. Although these regions are often considered motor output areas, numerous studies have shown that PMd, in particular, plays an active role in movement planning and decision formation prior to movement initiation (Confais *et al*., 2012; Nougaret *et al*., 2024; López-Galdo *et al*., 2025). Our results highlight three main observations regarding the relationship between timescales and task-related encoding in PMd and M1.

First, we found largely no differences in instantaneous decoding accuracy between short tau and long tau neuronal populations. This result contrasts with previous findings in PMd and other prefrontal structures where neurons with longer timescales were reported to carry stronger or more predictive task signals (Cavanagh *et al*., 2016, 2018; Cirillo *et al*., 2018; Fascianelli *et al*., 2019). Importantly, in our data, this discrepancy cannot be explained by insufficient availability of task information or by time window too limited in duration since our pre-Go delay was longer than the one reported by a previous study (Cirillo *et al*., 2018). These findings indicate that timescale does not necessarily determine the amount of information encoded at a given time point, even in conditions where task-relevant representations are robust and sustained. These two populations could play complementary roles within the population code that maintain information over time. Short-tau neurons display a transient selectivity and enable rapid encoding of sensory stimuli as reported in sensory areas (Murray *et al*., 2014). Long-tau neurons display a persistent activity over time with a more stable representation of information.

Second, during the last delay before the Go signal, PMd long-tau neurons showed a more stable representation of information. Indeed, functional relevance of timescales is about not only presence or absence of information but also how that information is structured in time. Long tau PMd neurons showed a more stable representation of direction information during the last delay before the Go signal than short tau neurons, indicating that the neural code for future movement is preserved over time during this period. This is consistent with population-level persistent coding within a stable representational subspace (Cavanagh *et al*., 2018; Cavanagh, Hunt and Kennerley, 2020) and with attractor-like dynamics that could support the maintenance information until action is required (Wang, 2001). Third, during movement execution, diagonals of the cross-temporal generalization decoding matrices were wider for long-tau neurons. Thus, during movement execution, both M1 and PMd displayed greater temporal stability of movement-direction encoding in long-tau neurons. This pattern suggests that long-tau neurons may support more sustained representations during the execution of the motor plan, whereas short-tau neurons may contribute to more rapid or intermittent updating of the evolving motor output.

## Methods

### Subjects

Two adult male Rhesus monkeys (*macaca mulatta*) (T and M, 10 and 14 kg, respectively) participated in this study. The inclusion of two monkeys follows standard practice in the field. The care and treatment of the animals conformed to the European Commission Regulations (Directive 2010/63/EU on the protection of animals used for scientific purposes) applied to French laws (decision of the 1st of February 2013), during all stages of the experiments. The experimental protocol was carried out in a licensed institution (B1301404) under the authorization 03383.02 delivered by the French Ministry of High Education and Research and evaluated by the local Ethics Committee (CEEA 071). The two macaques were monitored daily, by the animal care staff and the researchers involved in the study. The facility veterinary controlled regularly the general health and welfare conditions of the animals. The animals were pair-housed, and toys and enrichment, usually filled with treats, were routinely introduced in their home cage to promote exploratory behavior. During task performance, the animals received liquid reward from a dispenser. The animals were water-restricted in their home cage, with free access to dry pellets. In the event of reduced liquid consumption during task performance, the minimum daily intake was reached by giving extra water and fruit or vegetables in the home cage, delayed for a few hours after the end of the behavioral session. The daily fluid intake was never below 18 ml/kg, a low level for which it has been shown that macaques are able to effectively modulate their blood osmolality (Yamada, Louie and Glimcher, 2010), based on each animal’s reference body weight (measured prior to entering the liquid restriction regime). On resting days (e.g. weekends), the animals received a complete ration of liquid in the form of water and fruits in the home cage. Previous studies used data from these same two monkeys, recorded in the same hemisphere (Nougaret *et al*., 2024; López-Galdo *et al*., 2025), or from the other hemisphere (Kilavik, 2010; Ponce-Alvarez, Kilavik and Riehle, 2010; Confais *et al*., 2012, 2020; Kilavik *et al*., 2012; Kilavik, Confais and Riehle, 2014), during hand reaching behavior, and from the same hemisphere in monkey T in resting state sessions (Morales-Gregorio *et al*., 2025).

### Behavioral setup and task

A detailed description of the task design and data collection procedures can be found in a previous study (Nougaret *et al*., 2024). Two monkeys were trained to perform a visuomotor delayed-match-to- sample (DMS) task with fixed cue order and a Go signal, using their right arm. They performed arm-reaching movements in one of four diagonal directions from a central starting position. The monkeys controlled a freely movable handle on a horizontal plane while a visual display was presented on a vertical computer monitor (LCD; 75Hz) positioned in front of them, at 57 cm (Fig. 1A).

At the start of each trial, the display showed the handle’s position (small white square, 0.4 cm radius), a central fixation spot (yellow flashing disc, 0.45 cm radius), and four possible peripheral targets (red circular outlines, 1.5 cm radius) located 9 cm diagonally from the center. The monkey initiated a trial by positioning the cursor within the central fixation spot. Once the cursor was correctly placed, the fixation spot stopped flashing (though it remained visible), accompanied by a 50 ms tone. After maintaining the central hold for 1000 ms, a selection cue (SEL) appeared on the screen for 300 ms. The SEL was a large polygon (∼3 cm radius) in one of three colors (blue, green, or pink), indicating the color to attend to, for that trial. Following a 1000 ms delay, three spatial cues (SC1, SC2, SC3) were presented sequentially, each for 300 ms and each followed by a 1000 ms delay. These SCs were small colored discs (0.9 cm radius), always presented in the temporal order blue-green-pink, and each could appear in any of the four peripheral target locations (equally likely, with possible repetitions within a trial). This design yielded a total of 192 unique trial types, combining the three-color conditions with the four independent positions for SC1, SC2, and SC3. The monkey’s task was to remember the location of the SC matching the SEL color (valid cue) and ignore the two distractor SCs. After the final delay following SC3, a Go signal represented by simultaneous onset of the four red LEDs embedded in front of the monitor, prompted the animal to execute a center-out reach towards the memorized target location. The Go signal provided no directional information. Reaction time and movement time were each limited to 500 ms. Upon entering the correct target, the animal held position for 300 ms to receive a reward. Touching the valid target triggered a 50 ms tone, and a second tone marked the end of the hold period. A small drop of water or diluted fruit juice was delivered 500 ms later as a reward.

To simplify task demands, the three-color conditions were presented in separate small blocks of approximately 15 unique trials each, cycling through multiple blocks to cover all conditions. Trials within a block were presented in pseudo-random order, and any incorrect trial was re-presented later in the same block. A block was only considered complete when the monkey correctly performed all it’s unique trials. For monkey T, who completed fewer trials than monkey M, only three of the four target positions were used per session (randomly selected each time) to reduce the total number of unique trials. The monkey’s manual (horizontal) workspace was proportionally reduced relative to the visual display on the monitor, by a factor of approximately 0.7. In this physical workspace, the diagonal distance between the central fixation point and each peripheral target (center to center) was 6.5 cm. The allowable fixation zone was defined as a circle with a 0.3 cm radius, while the acceptable touch zone for peripheral targets had a radius of 1 cm. A response was considered valid when the hand-controlled cursor overlapped by more than half with the fixation or target outline.

In summary, the DMS task consisted of a predictable Go timing and sequential order of the three SCs, with SC validity indicated by the SEL cue at trial onset. The only unpredictable element was the spatial position of the three SCs.

### Data Acquisition

Recordings were conducted up to five days per week. On each recording day, a computer-controlled multielectrode microdrive (MT-EPS, Alpha Omega, Nazareth Illith, Israel) was mounted onto the recording chamber to transdurally insert electrodes into the motor cortex. Depending on the session, up to five single-tip microelectrodes (0.3–1.2 MΩ at 1 kHz; FHC) or up to two linear microelectrode arrays (V- or S-probes, Plexon, Dallas, TX, USA; or LMA arrays, Alpha Omega) were used. Arrays contained 24 or 32 contacts with 100, 150 or 200 μm inter-contact spacing and 12.5–15 μm contact diameters.

In this study, each recording from a single-tip electrode or one linear array within a behavioral session is referred to as a ‘site’. Electrodes and arrays were positioned and lowered independently using the Flex-MT drive (Alpha Omega). Individual guide tubes were used that did not penetrate the dura (no guide was needed for the rigid LMA arrays).

For single-tip electrodes, a common reference was used for all channels and connected together with the ground to a metal screw on the saline-filled titanium recording chamber. For linear arrays, the reference configuration depended on the array type. For LMAs (Alpha Omega), the reference was an insulated wire with its tip exposed either in the chamber saline or clipped to the stainless-steel probe tube (immersed in saline but insulated from brain tissue). For Plexon V-and S-probes, the reference was typically the stainless-steel probe shaft extending into the cortex.

In a few sessions, alternative references were used: for V-probes in monkey T, 6/36 sites were referenced to a posterior skull screw on the headpost; for S-probes in monkey M, 2/50 sites were referenced to a screw on the saline-filled chamber. For both array types, the ground was connected either to a skull screw on the titanium headpost or to a screw on the recording chamber.

Two data acquisition systems were used. All single-tip recordings in monkey T were obtained with an Alpha Omega platform comprising Alpha-Map software (Windows XP) and the MCP-Plus multichannel signal processor with analog headstages. Signals were amplified (gain 5,000–10,000), hardware filtered (1 Hz–10 kHz) and digitized at 32 kHz for offline analysis. All linear array recordings in monkey T, and all recordings in monkey M, were obtained using a Blackrock Neurotech system (Salt Lake City, UT, USA). This system employed Cereplex M digital headstages (PN 6956, 9360, 10129) connected via custom HDMI cables (PN 8083, 8068) to a Digital Hub (PN 6973, 6973 DEV 16–021, 10480), transmitting signals through fiber optics to a 128-channel Neural Signal Processor (NSP hardware version 1.0). Data was acquired with Cerebus Central Suite software (v6.03 for monkey T, v6.05 for monkey M; Windows 7). In monkey M, multiple single-tip electrodes were connected to the Cereplex M Omnetics connector via an adapter (PN 9038). Signals were hardware filtered (0.3 Hz–7.5 kHz) and digitized at 30 kHz for offline analysis.

Behavioral event codes (8-bit TTL) were transmitted in real time from the task control software (VCortex v2.2; NIMH, http://dally.nimh.nih.gov; Windows XP) to the DAQ system for synchronization with neural recordings.

### Recording sites

The monkeys were trained to perform the task using their right arm for multielectrode recordings from the left hemisphere (contralateral to the trained arm), targeting the dorsal premotor cortex (PMd) and the primary motor cortex (M1). Surgical preparation was performed in two stages. In a first surgery, a titanium head-post was implanted posteriorly on the skull and secured with titanium bone screws and bone cement. This was done after the monkeys had learned the basic visuomotor task. This allowed for head fixation during final training and recording sessions. Several months later, after the monkeys had fully learned the task and to work with head fixation, a second surgery was performed to implant a cylindrical titanium recording chamber (19 mm inner diameter) over the left hemisphere. The recording sites included in this study spanned approximately 15 mm along the anterior-posterior axis of the cortical surface. The approximate position of the chamber above M1 and PMd was confirmed with T1-weighted MRI scans (prior to surgery in both animals, and also post-mortem in monkey M). Intra-cortical electrical microstimulation (ICMS; (Asanuma and Rosén, 1972)) was used to identify and include only sites determined with ICMS to be related to upper limb movements.

### PMd/M1 classification

In this study, the classification of recording sites into PMd and M1 was based upon the dominant LFP beta-band power (Fig. 1B). We recently reported that low beta (<20 Hz) is strongest in M1 and relates to movement preparation and spontaneous posture, while high beta (>20Hz) is strongest in PMd and reflects temporal prediction and visuospatial attention (Nougaret *et al*., 2024). Beta power thus provides a functional split between the two motor cortical areas. For each recording site, we computed the periodic-only beta power in two frequency ranges 13-19Hz (low beta) and 23-29Hz (high beta) and determined whether low- or high-beta activity was dominant. This functional M1/PMd border was defined as the line that best separated the large majority of anterior sites with high beta dominance and the large majority of posterior sites with low dominance in each monkey (Nougaret *et al*., 2024).

### Identification of superficial (SUP) vs deep (DEEP) layers

For each recording site, linear probes were inserted gradually while monitoring neural activity to identify the depth at which the first neuronal signals appeared on the deepest probe contacts, both audibly and on the data acquisition monitor (DAQ). Electrodes immediately above them typically showed a prominent heartbeat-related signal in the LFP, confirming the insertion of the deepest probe contacts within the dura. More superficial contacts exhibited higher noise levels, including the 50 Hz line noise, as the recording chamber was filled with saline only after correct probe positioning. The probe was then lowered in small, regular steps of 50–100 μm, with neuronal and heartbeat signals shifting progressively upward across contacts. Once all but 2–3 contacts were within the brain, the probe was left in place for 1–3 hrs to allow tissue stabilization before recordings began. If neuronal activity was subsequently observed on all contacts, the probe was retracted until heartbeat signals were present on the top 1–2 contacts and neuronal activity was restricted to contacts 2 or 3 and below. In some cases, minor upward tissue drift occurred during recordings, but this typically did not exceed the inter-contact spacing.

A slightly tilted adapter mounted on the recording chamber ensured that probes entered the cortex approximately perpendicular to the cortical layers, provided recordings were conducted sufficiently far from major anatomical landmarks such as the central sulcus, arcuate sulcus, and precentral dimple. The locations of these landmarks were estimated from anatomical MRI scans and further verified using intracortical microstimulation (ICMS) chamber mapping with single electrodes during initial recording sessions.

Cortical layers were defined on a site-by-site basis according to depth relative to the dura. On the gyral surface, macaque M1 and PMd are approximately 3.5 mm thick (Koo *et al*., 2012; Beul and Hilgetag, 2019) also confirmed in our animals using anatomical MRI. Given that superficial and deep layers in macaque cortex are approximately equal in thickness (Hutsler, Lee and Porter, 2005), the upper 1.8 mm was classified as superficial layers (L2/3), with deeper contacts assigned to deep layers (L5/6). Because some linear arrays extended to depths of up to 4 mm, channels assigned to layer VI may have included interstitial neurons located within the underlying white matter. These neurons are thought to correspond to late-born layer VI cells that did not fully migrate into the cortical plate during development (Kostovic and Rakic, 1980). Given their similar morphological and functional characteristics, such interstitial neurons are commonly grouped together with layer VI neurons.

Laminar identification based on current source density (CSD) analysis, which is widely used in primary visual cortex to localize layer IV, is less reliable in higher-order cortical areas with less distinct laminar organization, such as motor and premotor cortex. This limitation is particularly pronounced for identifying the input layer, which lacks a clearly defined granular structure in these regions.

### Spike sorting to extract single unit activity (SUA)

We used the Plexon Offline Sorter (version 3.3.3 for monkey T and 4.6.0 for monkey M) for offline spike sorting. Continuous data were first high-pass filtered within the software using a 4-pole elliptic filter, with 250Hz cutoff frequency. After this, spike detection thresholds were manually set for each channel based on visual inspection of the signal to ensure reliable separation of spike waveforms from background noise. Detected spike waveforms were then subjected to automatic clustering using the built-in k-means scan algorithm. The resulting clusters were manually inspected and curated. Cluster quality was evaluated based on waveform shape and stability, separation in feature space, interspike interval (ISI) distributions to verify the presence of a refractory period, and autocorrelograms. Cross-correlograms were additionally examined to confirm independence between simultaneously detected units on the same electrode. For each unit, the signal-to-noise ratio (SNR) was computed from the spike waveforms, defined as the mean trough-to-peak voltage amplitude divided by twice the standard deviation of the entire signal (Hatsopoulos, 2004). Only well-isolated clusters, with a SNR >3, furthermore, exhibiting stable waveform morphology and a clear refractory period were classified as single units and included in timescale analyses. Due to small drifts of the laminar probe within the brain tissue during sessions, an individual assessment was done for each sorted unit to determine the period within the complete recording session in which the unit was sufficiently well isolated, and only this period was used for analysis. For further analysis, spike times were downsampled to 1 kHz.

### Estimation of timescales (tau/*τ*) from single neuron activity

Single neuron timescales (tau/*τ*) were obtained from spike count autocorrelogram (AC), based on the method proposed by (Fontanier *et al*., 2022). The original R code published in this study was adapted into Python to compute the AC curve, identify peaks, and fit the curve with an exponential decay function. Python’s in-built functions were used to replace the R modules for smoothing and curve fitting, as detailed below.

For each neuron, interspike intervals (ISIs) were computed between each spike and its subsequent spikes, up to the 100th order, from the full spiking activity recorded in each session. The ISIs were then sorted into small bins of 3.33ms bin width, ranging from 0 to 1000ms. To generate the autocorrelation curve, the ISI counts in each bin were normalized by dividing by the total number of counts and then dividing by bin width (3.33ms). The resulting AC was smoothed using the locally weighted scatterplot smoothing (LOESS) method with a frac parameter of 0.1 and the loess_1d algorithm (loess version 2.0.11; (Cappellari *et al*., 2013)). This smoothing step reduced high-frequency noise, facilitating accurate peak detection. ISI bins centered at less than 10 ms were excluded to prevent contamination from intervals below the absolute refractory period, and the bin counts were adjusted accordingly. The peak of the AC was identified as the global maximum, with the exception that if the first bin contained the maximum, it was disregarded in favor of the next highest peak. We then fit a mono-exponential decay function (GLOBAL fit) to the portion of the AC following this peak using the curve_fit function from scipy.optimize in Python, with the following equation : AC ∼ Ae-t/*τ*+B, where A (amplitude) and B (offset) are positive constants. Initial guesses for fitting were generated within specific ranges: [0, 1000] for *τ*, and random uniform distributions between [0, 2*(max(AC) – min(AC))] for A and [0, min(AC)] for B. Each AC was fitted 50 times with fit errors estimated for each iteration. To detect multiple peaks in the AC, termed ‘SLOW’ and ‘FAST’ fits as per (Fontanier et al., 2022), the presence of a ‘dip’ between the first peak and 100 ms post-peak was assessed. The local minimum within this range was identified, and if the difference between the global maximum and the local minimum exceeded 75% of the total difference between the maximum and minimum, it was classified as a true ‘dip’. To find the second peak, the AC was examined from 12 ms after the dip onward, and the highest point was marked as the second peak. Two exponential decay curves were then fitted: one from the first peak to the dip (‘FAST’ fit) and another from the second peak to the end (‘SLOW’ fit), using the same initial conditions and fitting procedure. In the post hoc analysis, only fits with positive A and B values, *τ* within 0–1000 ms, and finite standard deviation estimates of *τ* were retained. Among the 50 iterations per type (‘GLOBAL’, ‘SLOW’, and ‘FAST’), the best fit was selected based on the lowest fit error. In cases where all three fits were available, the ‘GLOBAL’ fit was preferred if: (1) its root mean square error (RMSE) was lower than the combined RMSE of the ‘SLOW’ and ‘FAST’ fits, or (2) one or both ‘SLOW’ and ‘FAST’ fits were invalid.

Neurons were included in laminar and cross-structural analyses only if they exhibited a valid “GLOBAL” fit with a τ standard deviation below 150 ms. After applying these criteria, the dataset comprised of 964 units (89%) from 33 laminar sessions in monkey M, 457 units (92.5%) from 32 laminar sessions in monkey T, and 48 units (89%) from 25 single-tip sessions in monkey T, utilized for all subsequent analyses.

### Estimation of timescales (tau/*τ*)in separate trial epochs

To examine whether timescales differed between specific task epochs, we repeated the timescale estimation procedure on trial-aligned spike trains. The epochs of interest were preSC1 (2 s before the first spatial cue, prior to direction information), preGo (2 s before Go signal onset, after direction information was received and maintained in memory), and the complete trial (touch to Go signal onset). Trials were aligned on the onset of the SEL cue and truncated to the corresponding 2 s pre-event window for preSC1 and preGo epochs. For the preSC1 epoch, all SEL cue trials were included, as movement direction was not yet specified. For the preGo epoch, only trials in which direction information had already been established were analyzed (blue and green SEL cue). This allowed comparison of timescales between an epoch lacking directional information (preSC1) and one in which direction was encoded and actively maintained in working memory (preGo). For each neuron, ISIs were computed separately for each trial and concatenated across all trials in the epoch before binning (3.33 ms width, 0-1000 ms range). The total number of ISIs in these lagged bins was typically lower than in the full-session analysis, reflecting the selectivity of individual neurons to stimuli in specific epochs. Given the reduced spike counts, LOESS smoothing was omitted, and the exponential decay fitting was performed only when bins with zero ISI counts made up less than 50% of all bins. All other steps, peak detection, fitting, and neuron selection, were identical to the full-session analysis. Neurons were included if they had a valid global τ with standard deviation < 150 ms.

### Preprocessing and PSD computation of bipolar LFP

The detailed preprocessing of the raw signals are mentioned in an earlier study (Nougaret *et al*., 2024). The raw neural signals were processed offline to extract the local field potential (LFP). First, a zero-phase 4th-order Butterworth filter with a 250 Hz cut-off frequency was applied (using the butter and filtfilt functions in MATLAB) to low-pass filter the data. The LFP was then high-pass filtered at 1Hz, and power-line noise was removed using a 4th-order Butterworth band-stop filter (49.5 - 50.5 Hz). The filtered signals were then downsampled to 1 kHz and saved for further analysis. To ensure data quality, trials containing noticeable artifacts, such as those caused by teeth grinding, static electricity, or heartbeat signals, were identified through visual inspection and excluded (12.3% of all trials in monkey T and 5.1% in monkey M).

LFPs represent the low-frequency components of neural signals recorded by electrodes, reflecting the summed activity of a population of neurons. Since LFPs are a population measure, they can often capture shared fluctuations across nearby electrodes, particularly in laminar probes where contacts are in proximity. Such shared modulations can arise from common synaptic inputs to the recorded area or from volume conduction effects, reflecting non-local neural activity. To isolate locally generated activity, we applied a bipolar derivation approach, a referencing method that estimates local signals by subtracting the potentials recorded at two adjacent electrodes referenced to a common ground. Since our recordings included probes with inter-contact spacings of 100, 150, or 200 µm, we first interpolated the signals from the 150 µm probes to obtain a uniform 100 µm depth resolution before computing bipolar derivations. For these interpolations, the signal at each intermediate depth was estimated as a weighted sum of the two adjacent channels, with weights determined by their relative distances. Finally, the ‘local’ bipolar signals were obtained by subtracting signals from electrode pairs separated by 400 µm. The bipolar signals were also aligned on ‘SEL’ (see Behavioral setup and task) before computing their PSDs. In total, monkey M contributed 31 laminar behavioral sessions, while monkey T contributed 23 laminar sessions. Across these sessions, 933 bipolar channels were included for LFP analyses in monkey M and 651 in monkey T. These channels were used for comparisons between PMd and M1 and, within each area, further categorized into superficial and deep layers.

PSD analysis of the aligned bipolar signals was performed using the conventional Welch’s method, implemented via the Python mne toolbox (version 1.7.1). This method computes short-time Fourier transforms from the time series and takes the mean across time to produce a smoothed estimate of the power spectrum. For our analysis, we used 1-second-long Hamming windows with 500 ms overlap between adjacent windows. The power spectra were computed over a frequency range of 0 to 100 Hz.

### Estimation of timescales from LFP Power spectra

In addition to traditional autocorrelation analysis of single-neuron activity, timescales have also been inferred from the power spectra of population signals such as LFPs (Manea *et al*., 2024) and ECoG (Gao *et al*., 2020). This study (Gao *et al*., 2020) demonstrated that neuronal timescales can be derived in the frequency domain since the power spectral density (PSD) is mathematically equivalent to the autocorrelation function (ACF) through the Wiener–Khinchin theorem (i.e., the PSD is the Fourier Transform of the ACF). Estimating τ in the frequency domain allows for simultaneous removal of oscillatory components (modeled as Gaussian peaks) and incorporation of variable power-law exponents, yielding a more accurate measure of the underlying aperiodic dynamics than directly fitting the ACF. In this framework, rather than computing the decay constant of the ACF, one identifies the knee frequency (*f_k_*), the frequency at which the power spectrum transitions from a flat low-frequency region to a 1/f-like decay at higher frequencies according to power-law (power ∝ 1/f ). From this knee frequency, neural timescale (decay constant *τ*) can be approximated as *τ* ≈ 1/2*πf_k_* , reflecting an inverse relationship between knee frequency and timescale: as *f_k_* increases, *τ* decreases.

To estimate the aperiodic component of the LFP spectra, we used the FOOOF algorithm (version 1.1.0; (Donoghue et al., 2020)) implemented in Python and applied to bipolar LFP signals. Power spectra were computed by averaging across trials in the 2–80 Hz range. The algorithm was run in knee mode (peak_width_limits = [2, 12]) to estimate both the spectral exponent ‘*χ*’ (exp) and the knee parameter ‘k’. The corresponding knee frequency (*f_k_*) was computed as *f_k_* = *k*^1/^*χ*. However, only 23% of channels across both monkeys and recording sites exhibited a reliable knee parameter. Therefore, to obtain timescale estimates across the full dataset, we quantified temporal autocorrelation of the bipolar LFP signals and estimated the aperiodic spectral exponent as additional measures of local circuit dynamics.

### Estimation of timescales from autocorrelation of LFP signals

Timescales were estimated from bipolar signals using the Python-based toolbox timescales-methods (Voytek Lab, [timescales toolbox](https://voyteklab.com/timescale-methods/index.html)). The method models the aperiodic component of the autocorrelation function (ACF) using an exponential decay function and extracts the decay constant (*τ*/tau) as the timescale estimate. To ensure that the estimated tau reflected background neural dynamics rather than oscillatory structure, the fitting procedure was performed with the parameter with_cos = False. This prevented the model from fitting periodic components (the peaks and trough) of the ACF. To ensure fit quality at single-trial level, a minimum *R*^2^ threshold was kept at 0.35. Trials with fits below this threshold were excluded. Finally, the median tau across retained trials was computed and used as the representative LFP timescale for each bipolar channel within a recording session.

### Estimation of aperiodic exponents using FOOOF

The spectral exponent (‘*χ*’ or exp) represents the slope of the aperiodic component in log-log power spectra and captures how power decreases with increasing frequency. Previous work (Gao et al., 2017) demonstrated that the exponent reflects the underlying excitation-inhibition (E/I) balance, particularly within the 30-70 Hz range of LFP power spectra. More recently, (Cusinato *et al*., 2023) showed that spectral exponents follow a hierarchical gradient across brain regions, from temporal lobe to hippocampal to amygdalar areas, which inversely mirrored the hierarchy of timescales observed in these areas.

In our dataset, approximately 23% of channels exhibited a knee in the aperiodic component, with the max knee frequency observed at 39 Hz. To obtain comparable exponent estimates across all channels while minimizing contamination from beta oscillations (13-30Hz) and avoiding the knee region, we restricted the fitting range to 55-95 Hz. The aperiodic exponent was extracted by fitting the average power spectral density (PSD) across trials using the FOOOF toolbox, with peak_width_limits = [2.0, 12.0] and mode = ’fixed’. Thus we obtained spectral decay estimated for each bipolar channel in our dataset.

### Channel inclusion for estimating spectral exponents

All analyses involving spectral exponents were performed using values derived from the full-session bipolar channels. Since bipolar channels are constructed by differencing pairs of unipolar electrodes, some bipolar channels span different cortical layers, specifically, those combining superficial and deep contacts. To ensure laminar specificity, these mixed-layer bipolar channels were excluded from laminar analyses but retained for comparisons at the regional level between PMd and M1. Detailed counts of unipolar and bipolar channels used across all analyses, including distinctions between laminar and region-level comparisons, are provided in Table 1 and Supplementary Table 1.

### Statistical test

Statistical comparisons of timescales (tau) and aperiodic exponents (exp) across areas (PMd vs M1) and layers (superficial vs deep) were performed using non-parametric Kruskal–Wallis tests, with significance indicated as follows: “* p < 0.05, ** p < 0.005, *** p < 0.0005.”

### Decoding analysis of movement direction

To evaluate the link between intrinsic neuronal properties (tau) and their involvement on cognitive and motor processes, we used a decoding procedure to discriminate movement direction along task epochs. Monkey T was performing the task with only 3 different directions in a given recording session, differently than monkey M performing the task with all 4 directions in each session. Consequently, to pool data from two monkeys, we first reassign direction labels of Monkey to have unique labels of ‘remapped direction’ 1, 2 and 3 that correspond to the 3 directions used in that session, losing the ‘real direction’ labels (up-left / up-right / bottom-right / bottom-left). Then, we selected for each session of Monkey M, the 3 directions in which the monkey performed the highest number of correct trials and assigned to them the ‘remapped direction’. We used the Neural Decoding Toolbox (Meyers, 2013) for all decoding analysis. For each neuron, data were binned in the epoch of interest (50ms bins without overlap from -1200 ms to +7500 ms from SEL appearance or from -1000 ms to + 2000ms from hand movement onset). For each bin, a different classifier was trained and tested. A z-score normalization was applied to each neuron across trials to give equal weight to all recorded neurons regardless of their firing rates. Trials were labelled depending on the condition (remapped direction 1, 2 or 3) and the maximum number of trials, *k*, available by condition for the maximal number of neurons was estimated. A trade-off between excluding the smallest number of neurons and keeping the maximum number of trials led to *k* = 8 when the analysis were targeting only one color condition (blue, green or pink conditions, all alignments on SEL appearance), and to *k* = 12 when all color conditions were combined (all alignments on hand movement onset). The classifier was trained using the following cross-validation procedure: trained on *k-1* splits and tested on the remaining split. The number of selected neurons for each decoding procedure is reported in Supplementary Table 2. This process was repeated 100 times, or runs, selecting randomly different subset of trials every time. The *real* classification accuracy reported in the figures is the average of these 100 runs. To assess the over chance significance of the decoding accuracy in each temporal bin, we also perform the same decoding procedure 100 times but with the labels randomly shuffled to obtain a null distribution of the shuffle data. To save computational time, each *shuffle* procedure was repeated only for one run. Statistical significance was assessed using a Monte-Carlo approximation of the p-value. For each time bin, the real decoding accuracy was compared against 100 shuffles. The p-value was calculated as: 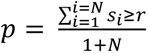 where r represents the real statistic, *s_i_* represents the shuffled statistic and N the number of shuffles.

To assess the significance between two *real* accuracies (i. e. short and long tau subpopulations of neurons), a one-sided empirical percentile criterion was used: at each time bin, the mean accuracy of the higher-accuracy condition had to exceed the 97.5th percentile of the lower-accuracy condition distribution, allowing effects in either direction. Two distributions were considered as significantly different for a percentile criterion reached for at least three consecutive time bins.

The cross-temporal decoding (CTD) maps were built by using the same decoding procedure described above but for all train/test bin possible combinations. A *real* decoding map was built by averaging maps from 100 runs and 100 *shuffle* maps were built with the labels randomly shuffled. The significance of each pixel of the map was assessed with a comparable percentile criterion than described above: the mean real accuracy had to exceed the 95th percentile of the shuffle accuracy condition distribution. The number of significant pixels between two maps were then compared using a permutation test on the proportion of significant pixels by comparing the observed difference to a null distribution obtained by randomly relabelling pixels. P-values were estimated using a Monte-Carlo approximation : 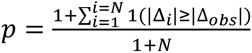 , where Δ*_obs_* is the observed difference in proportions Δ*i* is the difference in proportions obtained at permutation i, N is the number of permutations (10000). The summation counts the number of permutations for which the absolute difference in proportions was at least as large as the observed difference.

Finally, for the analysis performed aligned on the hand movement onset, we computed the width of the cross-temporal decoding map diagonal. For train times from 0 to 500ms after hand movement onset, we realigned horizontal stripes of the CTD maps on the diagonal pixel and extracted a width profile of the diagonal. The full width at half-maximum (FWHM) of the diagonal decoding profile was computed relative to chance level and using linear interpolation to estimate the lag values at which the decoding curve crossed its half-maximum level. The FWHM was the difference between the interpolated right and left crossing points. Statistical significance of FWHM differences between two populations was assessed using a permutation test. Train-times profiles were pooled and randomly reassigned across conditions. For each permutation, mean profiles were recomputed, their FWHM estimated, and the width difference was calculated. P-values were estimated using the same Monte-Carlo approximation then previously described for proportion of significant pixels comparison.

## Acknowledgements

The authors thank Joel Baurberg, Xavier Degiovanni, and Luc Renaud for technical assistance; Sébastien Barniaud, Laurence Boes, Frédéric Charlin, and Marc Martin for animal care.

## Author Contributions

**NN:** Conceptualization, Methodology, Software, Formal analysis, Visualization, Writing - original draft, Writing – review & editing

**LLG**: Software, Visualization, Writing - review & editing.

**SN**: Conceptualization, Methodology, Software, Formal analysis, Supervision, Writing - review & editing.

**BEK**: Conceptualization, Software, Methodology, Resources, Data Curation, Investigation, Supervision, Writing - review & editing, Funding acquisition.

---

**Supplementary Fig 1.**
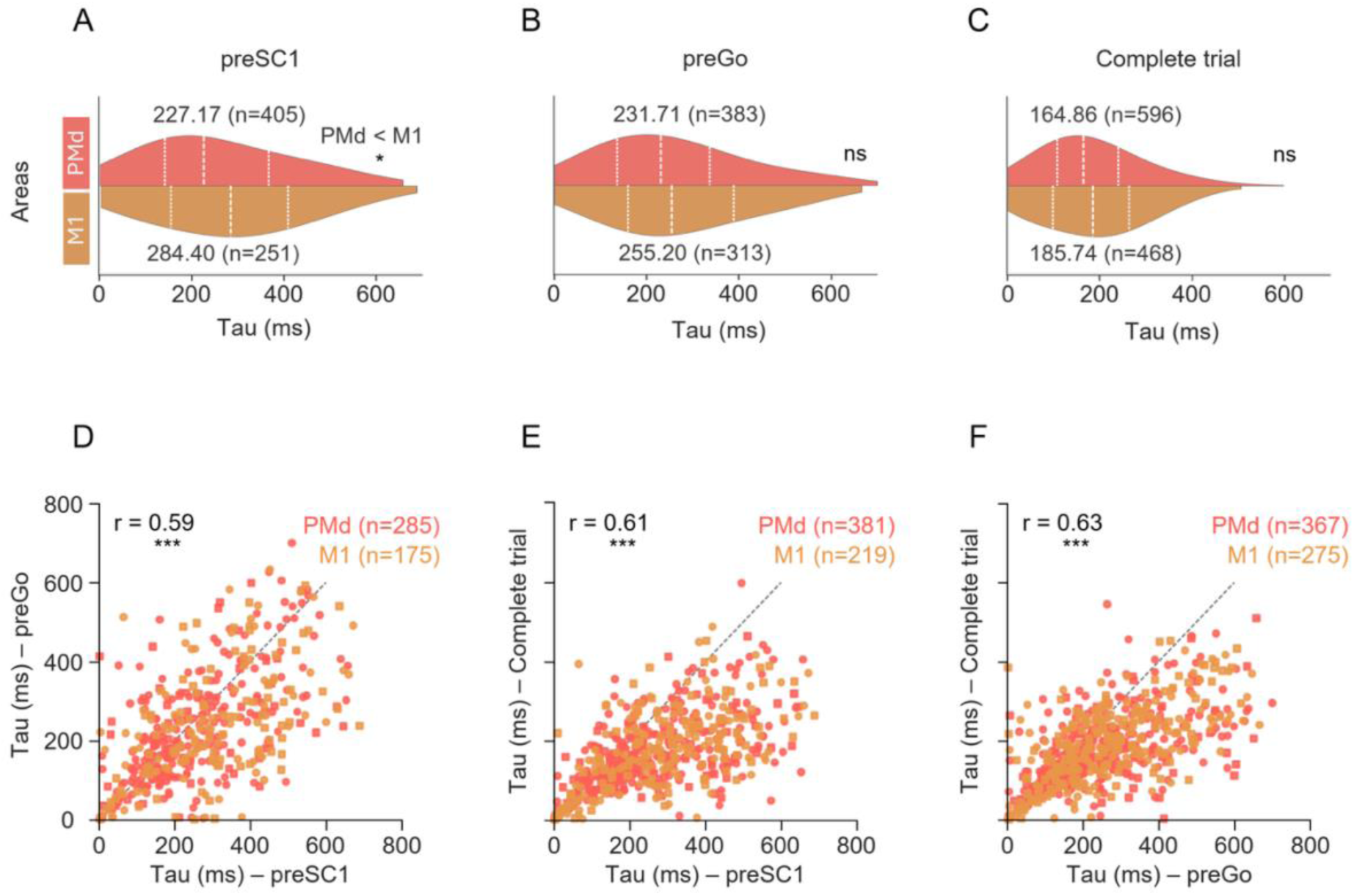
Comparison of SUA timescales across task epochs and areas (A-C) Spike tau (Tau) distributions for PMd and M1 neurons estimated from three separate task epochs: **(A)** preSC1, a 2-second window prior to the first spatial cue onset, before directional information was presented, **(B)** preGo, a 2-second window preceding the Go signal, after directional information was presented and **(C)** Complete trial epoch from touching the central fixation circle to the onset of Go signal (touch to Go). Dashed lines indicate the median, 1^st^ and 3^rd^ quartiles tau values for each area. Numbers indicate median values, and n denotes the number of neurons yielding valid tau estimates within each epoch. A significant difference between areas was observed only in the preSC1 epoch (*p = 0.01), and a similar trend in preGo and complete trial epochs, with longer tau in M1. **(D-F)** Spearman rank correlation of spike-tau estimates across pairs of task epochs: **(D)** preSC1 vs preGo, **(E)** preSC1 vs Complete trial, and **(F)** preGo vs Complete trial. Each data point represents one neuron, with circles corresponding to monkey M and squares to monkey T. n indicates the number of neurons with valid tau estimates in both epochs being compared. Correlations were significant across all epoch pairs (***p < 0.0001), indicating that spike-derived tau are largely preserved across distinct task periods.

**Supplementary Fig 2.**
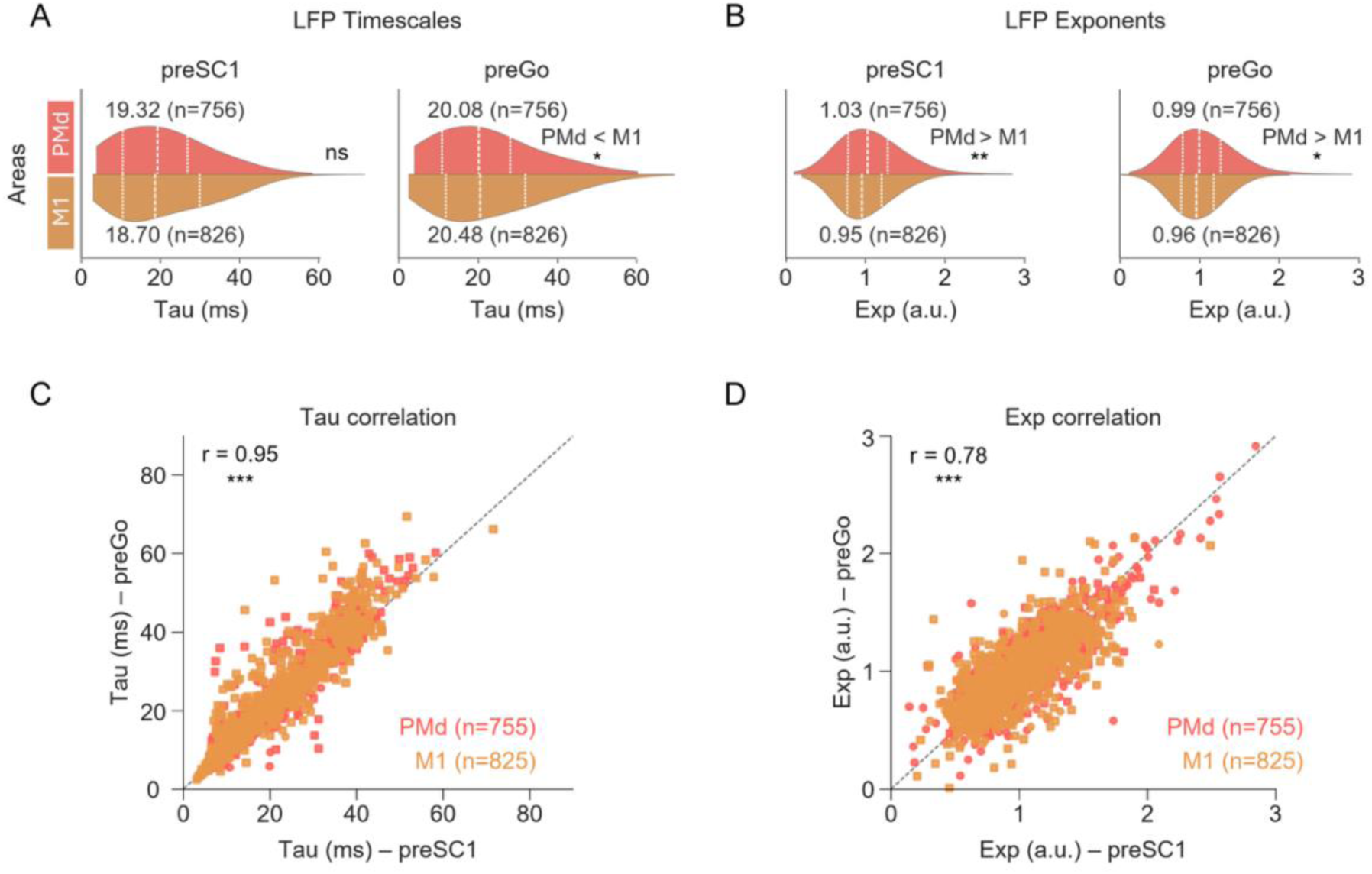
Comparison of LFP temporal measures across task epochs and areas (A–B) Distributions of LFP-derived timescales **(tau; A)** and LFP aperiodic spectral exponents **(exp; B).** The left subplot shows the measures (tau or exp) estimated during the preSC1 epoch, a 2-s window preceding the first spatial cue onset, before directional information was presented. The right subplot shows the estimation in the preGo epoch, a 2-s window preceding the Go signal after directional information had been presented. Dashed lines indicate the median, 1^st^ and 3^rd^ quartiles of each distribution. Numbers denote median values, and n indicates the number of bipolar channels yielding valid estimates within each epoch for each measure. For tau (A - right), a significant difference between PMd and M1 was observed only during the preGo epoch (p = 0.035), with longer timescales in M1. For exp (B), the PMd–M1 difference in exponents was significant in both epochs with PMd > M1 (preSC1 **p = 0.001 and preGo *p = 0.005). **(C-D)** Spearman rank correlation of LFP **(tau; C)** and spectral exponents **(exp; D)** estimated during the same preSC1 and preGo epochs. n indicates the number of bipolar channels with valid estimates for each measure (tau or exp) across both epochs. Correlations were significant for both tau and exp (***p < 0.0001), indicating that relative magnitudes of the LFP-derived temporal measures are preserved across distinct task epochs.

**Supplementary Fig 3.**
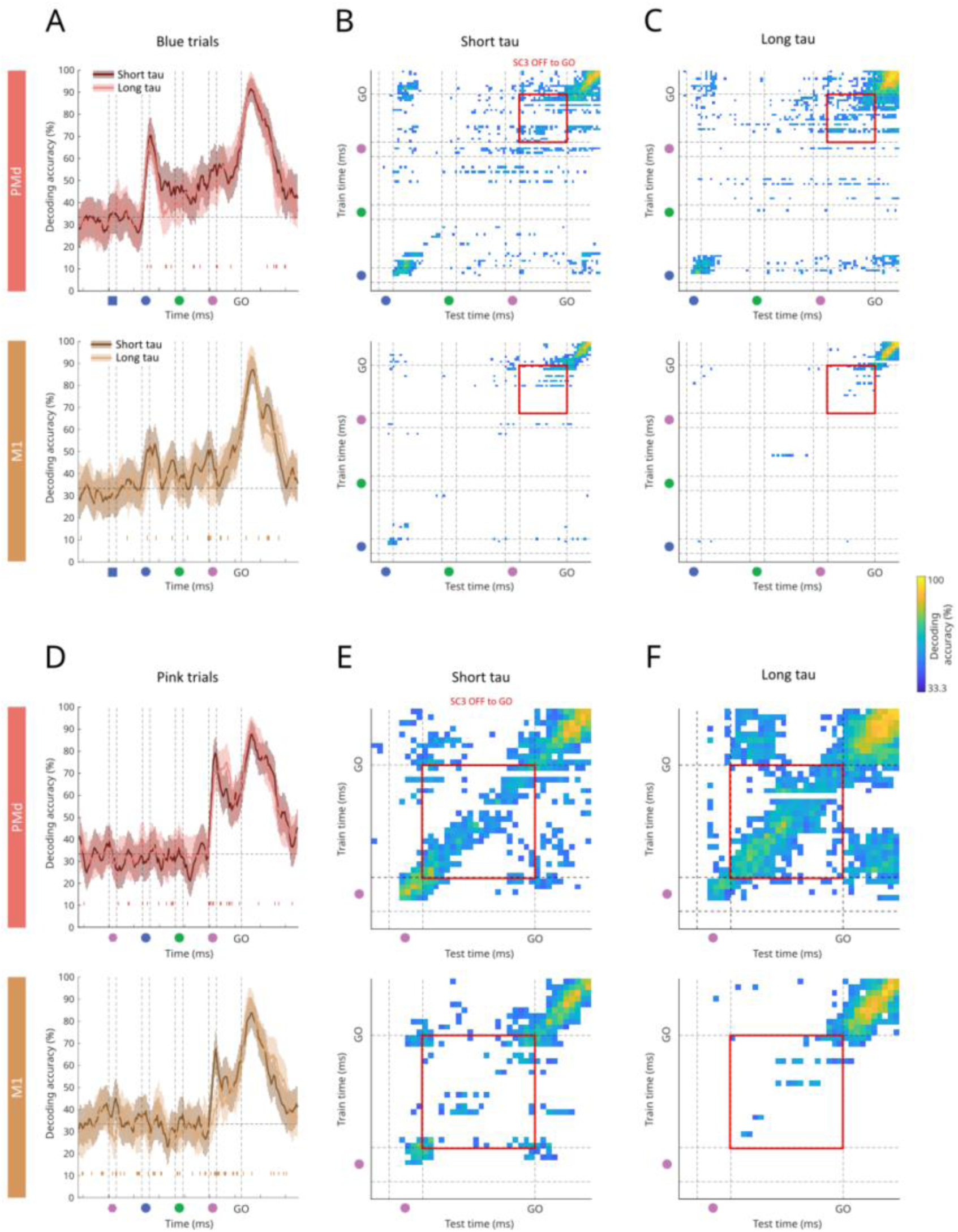
Direction encoding in short and long tau neurons during movement planning in blue and pink trials. Same representation as in (Fig 3B, C ,D). **(A, D)** Time-resolved classification accuracy (mean ± SD), for blue (A) and pink trials (D), for decoding performance of two classifiers built with either the 50% of short or long tau neurons subpopulations. **(B, E)** Cross-temporal decoding maps for short-tau neuronal population, in blue (C) and pink trials (E). **(C, F)** Same as in B, E, for long tau neuronal population. Permutation tests comparing proportions showed the following results: Blue trials PMd: 124/400 [31.00%] vs 144/400 [36.00%], p = 0.16; Pink trials PMd; 146/400 [36.5%] vs 267/400 [56.75%], p < 0.0001 ; Blue trials M1: 49/400 [12.25%] vs 32/400 [8.00%], p = 0.06 ; Pink trials M1: 60/400 [15.00%] vs 28/400 [7.00%], p < 0.001.

**Supplementary Fig 4.**
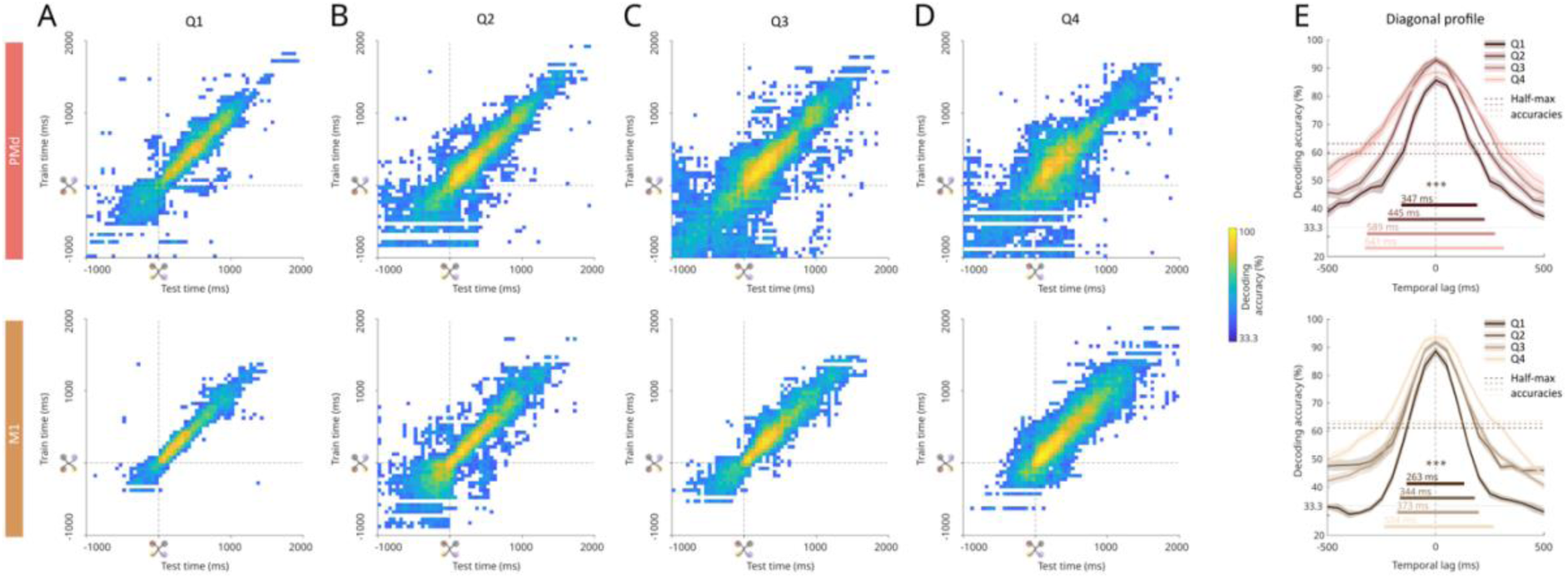
Direction encoding in PMd and M1 neurons across tau quartiles during movement execution. Same representation as in Figure 4B, C, D. **(A-D)** Cross-temporal decoding maps for Q1, Q2, Q3 and Q4 neuronal population, from -1000ms to +2000ms around movement onset. **(E)** Diagonal profile and half-max accuracies averaged across the first 500ms after movement onset. ***, permutation test between Q1 and Q4, p<0.001.

**Supplementary table 1.** Statistical comparisons of tau/exp (reported median values) in monkey M & T.

|  | Monkey T |  |  | Monkey M |  |  |
| --- | --- | --- | --- | --- | --- | --- |
|  | Comparison by Area / Layer |  | Kruskal Wallis | Comparison by Area / Layer |  | Kruskal Wallis |
|  | PMd | M1 |  | PMd | M1 |  |
| **Spike tau** | 182.57ms<br>(n=185) | 204.03ms<br>(n=320) | $\chi^2=1.74$,<br>$p=0.19$ | 140.67ms<br>(n=494) | 159.71ms<br>(n=470) | $\chi^2=6.49$,<br>$p=0.01$ |
| **LFP tau** | 31.58ms<br>(n=249) | 28.45ms<br>(n=401) | $\chi^2=9.3$,<br>$p=0.002$ | 17.74ms<br>(n=508) | 16.11ms<br>(n=425) | $\chi^2=2.75$, $p=0.1$ |
| **LFP exp** | 1.19<br>(n=249) | 1.07<br>(n=401) | $\chi^2=3.15$,<br>$p=0.08$ | 0.91<br>(n=508) | 0.87<br>(n=425) | $\chi^2=17.33$,<br>$p=3.14e-5$ |
|  | **PMd SUP** | **PMd DEEP** |  | **PMd SUP** | **PMd DEEP** |  |
| **Spike tau** | 165.89ms<br>(n=58) | 170.64ms<br>(n=80) | $\chi^2 = 0.3$, $p = 0.6$ | 146.12ms<br>(n=204) | 135.5ms<br>(n=278) | $\chi^2=0.85$,<br>$p=0.35$ |
| **LFP tau** | 30.91ms<br>(n=103) | 33.88ms<br>(n=90) | $\chi^2=0.16$, $p=0.7$ | 16.69 ms<br>(n=174) | 18.55 ms<br>(n=234) | $\chi^2=5.82$,<br>$p=0.02$ |
| **LFP exp** | 1.23<br>(n=103) | 1.18 (n=90) | $\chi^2=0.4$, $p=0.5$ | 0.98<br>(n=174) | 0.87<br>(n=234) | $\chi^2=12.52$,<br>$p=0.0004$ |
|  | **M1 SUP** | **M1 DEEP** |  | **M1 SUP** | **M1 DEEP** |  |
| **Spike tau** | 208.4ms<br>(n=158) | 167.3ms<br>(n=93) | $\chi^2 = 2.54$, $p = 0.1$ | 139.79ms<br>(n=169) | 163.37ms<br>(n=294) | $\chi^2=8.7$,<br>$p=0.003$ |
| **LFP tau** | 20.3 ms<br>(n=217) | 36.98ms<br>(n=96) | $\chi^2=66.6$,<br>$p=3.34e-16$ | 11.96 ms<br>(n=115) | 17.03 ms<br>(n=124) | $\chi^2= 25.18$,<br>$p=5.21e-7$ |
| **LFP exp** | 1.14<br>(n=217) | 0.83 (n=96) | $\chi^2=45.5$,<br>p=1.56e-11 | 0.92<br>(n=115) | 0.92<br>(n=124) | $\chi^2=0.001$,<br>p=0.97 |

**Supplementary table 2.** Number of selected neurons for each decoding procedure.

| Movement planning |  |  | Movement execution |  |  |
| --- | --- | --- | --- | --- | --- |
| Condition | PMd | M1 | Condition | PMd | M1 |
| **All neurons - blue** | 275/679 | 263/679 | Short tau - All colors | 264/333 | 284/395 |
| **All neurons - green** | 245/679 | 219/790 | Long tau - All colors | 258/331 | 280/395 |
| **All neurons - pink** | 232/679 | 204/790 | Q1 - All colors | 143/168 | 143/198 |
| **Short tau - blue** | 142/340 | 129/395 | Q2 - All colors | 121/165 | 141/197 |
| **Short tau - green** | 126/340 | 110/395 | Q3 - All colors | 124/164 | 139/198 |
| **Short tau - pink** | 120/340 | 106/395 | Q4 - All colors | 134/167 | 141/197 |
| **Long tau - blue** | 133/339 | 134/395 |  |  |  |
| **Long tau - green** | 119/339 | 109/395 |  |  |  |
| **Long tau - pink** | 112/339 | 98/395 |  |  |  |

